# Muskelin acts as a substrate receptor of the highly regulated *Drosophila* CTLH E3 ligase during the maternal-to-zygotic transition

**DOI:** 10.1101/2024.06.28.601265

**Authors:** Chloe A. Briney, Jesslyn C. Henriksen, Chenwei Lin, Lisa A. Jones, Leif Benner, Addison B. Rains, Roxana Gutierrez, Philip R. Gafken, Olivia S. Rissland

## Abstract

The maternal-to-zygotic transition (MZT) is a conserved developmental process where the maternally-derived protein and mRNA cache is replaced with newly made zygotic gene products. We have previously shown that in *Drosophila* the deposited RNA-binding proteins ME31B, Cup, and Trailer Hitch (TRAL) are ubiquitylated by the CTLH E3 ligase and cleared. However, the organization and regulation of the CTLH complex remain poorly understood in flies. In particular, *Drosophila* lacks an identifiable substrate adaptor, and the mechanisms restricting degradation of ME31B and its cofactors to the MZT are unknown. Here, we show that the developmental specificity of the CTLH complex is mediated by multi-pronged regulation, including transcriptional control by the transcription factor OVO and autoinhibition of the E3 ligase. One major regulatory target is the subunit Muskelin, which we demonstrate acts as a substrate adaptor for the *Drosophila* CTLH complex. Although conserved, Muskelin has structural roles in other species, suggesting a surprising functional plasticity. Finally, we find that Muskelin has few targets beyond the three known RNA binding proteins, showing exquisite target specificity. Thus, multiple levels of integrated regulation restrict the activity of the embryonic CTLH complex to early embryogenesis, seemingly with the goal of regulating three important RNA binding proteins.

## INTRODUCTION

The maternal-to-zygotic transition (MZT) is an essential process in early animal embryogenesis where the embryo switches from a maternally-deposited mRNA and protein cache to a zygotically-derived proteome and transcriptome. In mammals, the MZT occurs over the first 36-48 hours of development, while in *Drosophila melanogaster* the transition occurs in the syncytial zygote over the course of approximately 5 hours (Vastenhouw et al. 2019). Despite timescales being vastly different between different animals, the molecular cascades that orchestrate the MZT are remarkably well-conserved. Maternally deposited RNAs are rapidly destroyed in flies, zebrafish, mice, and humans (Bashirullah et al. 1999; Svoboda et al. 2015; Sha et al. 2020), and at the same time, zygotic transcription is initiated (Lee et al. 2013; Higuchi et al. 2018; Heyn et al. 2014; Duan et al. 2021; Harrison et al. 2011; Larson et al. 2022; Schulz and Harrison 2019). Although RNA regulation is much better understood than protein regulation in the MZT, a number of recent studies have highlighted the essential contribution of maternal protein decay during early embryogenesis (Zavortink et al. 2020; Cao et al. 2020, 2021; Gao et al. 2017; Dang et al. 2023; Kinterova et al. 2019; Shimuta et al. 2002).

One mechanism by which proteins are cleared is the ubiquitin-proteasome system (UPS). Here, a cascade of enzymes works together to create polyubiquitin chains on a target protein, marking it for degradation by the proteasome. First, an E1 ubiquitin-activating enzyme transfers a ubiquitin molecule to an E2 ubiquitin-conjugating enzyme, which then associates with an E3 ubiquitin ligase and the target, thereby catalyzing the ubiquitin transfer (Komander, David; Rape, Michael 2012). Target specificity is mainly orchestrated by the E3, which contains a substrate receptor domain or adaptor (Cowan and Ciulli 2022). One prototypical example is the family of RING-type E3 ligases, which catalyze ubiquitin transfer directly from the E2 to the substrate by correctly scaffolding the substrate, catalytic modules, and other supporting members of the complex (Deshaies and Joazeiro 2009; Rotin and Kumar 2009).

The CTLH E3 ligase is a multi-subunit, RING-type ligase that has emerged as a vital player in the *Drosophila* MZT. Known as the GID complex in yeast, this E3 ligase is conserved from yeast to humans and has been implicated in several important processes including early embryogenesis, erythropoiesis, immune response, tumorigenesis, and intermediary metabolism (Zavortink et al. 2020; Sherpa et al. 2022; Simwela et al. 2024; McTavish et al. 2019; Salemi et al. 2017; Liu et al. 2020; Santt et al. 2008; Qiao et al. 2020; Sherpa et al. 2021; Gottemukkala et al. 2024; Yi et al. 2024). In flies, the CTLH complex targets the RNA-binding protein ME31B and its two partners, Cup and Trailer Hitch (TRAL), which together form a post-transcriptional repressive complex and are critical for oogenesis (Wang et al. 2017; Tritschler et al. 2008; Zavortink et al. 2020; Wilhelm et al. 2005, 2003; Nakamura et al. 2001, 2004). All three proteins are maternally deposited in the embryo and then rapidly removed during the MZT by the ubiquitin-proteasome system via the CTLH complex (Wang et al. 2017; Zavortink et al. 2020; Cao et al. 2020). The degradation of ME31B and its partners is developmentally controlled, in part through a translational feedback loop: during oogenesis, the ME31B complex translationally represses the mRNA encoding the E2 dedicated to the CTLH complex (known as *Marie Kondo* or *Kdo*). However, this repression is alleviated upon egg activation through the action of the Pan Gu kinase, leading to Kdo production and full activation of the E3 ligase (Zavortink et al. 2020).

Much of the organization of the *Drosophila* CTLH complex has been inferred from work in yeast and humans (Figure 1A; Qiao et al. 2020; Sherpa et al. 2021; Maitland et al. 2022). Broadly, the E3 ligase has three organizational domains: catalytic, scaffolding, and substrate recognition. The catalytic components form a dimer and interact with its E2 via a set of conserved interactions (Chrustowicz et al. 2024). In yeast, a set of scaffold components, including Gid8 (known as Houki [Hou] in *Drosophila,* see Figure S1A for all homologous naming) and Gid1 (RanBPM in *Drosophila*), connect the catalytic domain to the substrate adaptor. The substrate recognition domain is composed of Gid4 and Gid5, which help identify and orient the target protein. In both yeast and humans, the complex is then organized into a higher-order supramolecular structure through an additional β-propeller subunit (Gid7 in yeast) that increases target affinity and ubiquitylation (Langlois et al. 2022; Sherpa et al. 2021). Although most subunits are broadly conserved with easily identifiable orthologs, *Drosophila* notably lacks the two subunits needed for substrate recognition in yeast: Gid4 and Gid5 (Figure 1A, Figure S1A; Zavortink et al. 2020).

**Figure 1.**
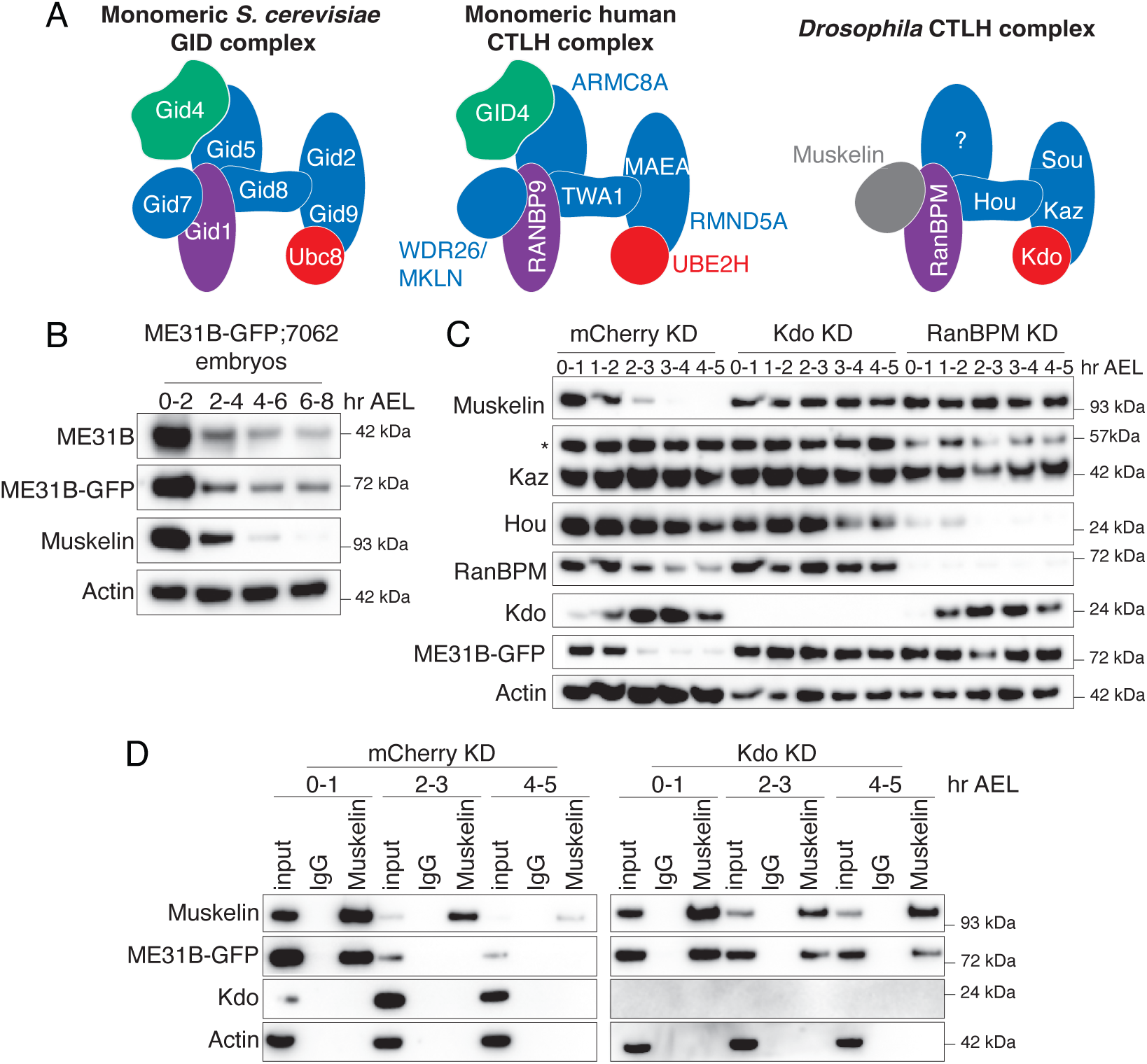
The embryonic CTLH complex is autoregulated and degraded at the end of the maternal-to-zygotic transition. A. Models of the CTLH complex in yeast, human, and fly. Substrate recognition module in green, catalytic/scaffolding modules in blue, and E2 in red. B. ME31B levels are stable after the end of the MZT. Lysates from staged embryos were probed for ME31B-GFP, ME31B, Muskelin, and actin (as a loading control) for the indicated time points. C. The CTLH complex is autoregulated. Staged embryos from the indicated times were harvested with mCherry, Kdo, or RanBPM depleted. Lysates were probed for the indicated proteins. (*) denotes non-specific band. D. The interaction between Muskelin and ME31B-GFP is more robust at the end of the MZT in the absence of Kdo. Muskelin was immunoprecipitated from staged lysates across the MZT time course from control (mCherry) knockdown or Kdo knockdown embryos. “Hr AEL”: hours after egg laying. All western blots are representative images from three biological replicates.

Target recognition is an essential aspect of how protein degradation by E3s, including the CTLH complex, is regulated. For instance, in yeast, the CTLH complex utilizes separate substrate adaptors, such as Gid4, Gid10, or Gid11, depending on the cellular environment and needs of the organism (Qiao et al. 2020; Gottemukkala et al. 2024; Sherpa et al. 2022; Maitland et al. 2024; Langlois et al. 2022; Kong et al. 2021). The situation is even more complex in humans, where different structural components, specifically the human orthologs of Gid7 (WDR26 and Muskelin), not only change the overall organization of the complex but also change substrate specificity. In particular, WDR26 plays a dual role in the human CTLH complex, functioning both in forming the supramolecular complex *and* in directly recognizing target proteins (Sherpa et al. 2021; Mohamed et al. 2021; Gross et al. 2024; Onea et al. 2022; Gottemukkala et al. 2024). Muskelin, the other Gid7 ortholog, is mutually exclusive with WDR26 and intriguingly, it is both ubiquitylated as part of autoregulation by the CTLH complex and regulates a distinct subset of the proteome compared to WDR26 (Maitland et al. 2019, 2024; Yi et al. 2024). Recent work has also demonstrated that Muskelin-mediated CTLH supramolecular complex formation is required for degradation of a metabolic enzyme in humans in response to mTORC1 inhibition (Yi et al. 2024; Maitland et al. 2024), although how Muskelin affects target specificity is unknown. Intriguingly, the *Drosophila* genome lacks orthologs for the best described substrate adaptor (*Gid4*) and even lacks an ortholog for *Gid5* (Zavortink et al. 2020). Moreover, the ortholog of *WDR26* in *Drosophila* (*CG7611*) is not required for the destruction of ME31B and its cofactors during the MZT (Zavortink et al. 2020), leaving open the question of the identity of the substrate receptor for the embryonic CTLH complex. In other words, two mysteries surround the *Drosophila* CTLH complex, particularly in the context of protein degradation during the MZT. First, its target ME31B is an essential RNA-binding protein that is broadly expressed alongside nearly all components of the CTLH complex: it has therefore been unclear what mechanisms might restrict the degradation of ME31B to the MZT. Given the notable absence of a *Gid4* ortholog, we reasoned that this mystery might be linked to a second one – the identity of the *Drosophila* CTLH substrate receptor.

Here, we describe multiple mechanisms that restrict the recognition of ME31B by the CTLH complex to the MZT beyond the previously-established translational control of *Kdo*. First, at the end of the MZT, the interaction between the CTLH complex and its target proteins weakens during the MZT, and then autoregulation further reduces levels of complex components. Third, transcription of the Muskelin subunit (which is required for ME31B degradation) is tightly restricted to oogenesis. Muskelin expression is mediated by the OVO oogenesis-specific transcription factor, thus providing an explanation for why ME31B is not targeted outside the MZT. Importantly, we also demonstrate that Muskelin functions as the substrate adaptor for the CTLH complex that recognizes ME31B. We found that the embryonic CTLH complex robustly targets, at most, four proteins during the MZT; in other words, ME31B and its binding partners appear to be the critical targets of Muskelin. The complex layers of transcriptional, post-transcriptional, and post-translational regulation thus converge to restrict the activity of the embryonic CTLH complex — and the degradation of ME31B/Cup/TRAL — to a tight developmental window.

## RESULTS

### The embryonic CTLH complex is autoregulated and degraded at the end of the maternal-to-zygotic transition

ME31B degradation in the MZT is mediated by the E2-E3 combination of *Marie Kondo* (*Kdo*) and the CTLH complex (Zavortink et al. 2020; Cao et al. 2020), but little is known about its degradation beyond the MZT. To investigate the temporal control of ME31B degradation, we performed an extended time course by harvesting protein lysate from embryos through and beyond the end of the MZT. As expected, both endogenous ME31B protein levels and overexpressed ME31B-GFP protein levels dropped quickly during the MZT (2-4 hours after egg laying), but they remained steady at the end of the MZT and beyond (Figure 1B).

We next developed antibodies to components of the CTLH complex (Muskelin, RanBPM, Katazuke [Kaz], and Houki [Hou]; Figure S1B) and probed levels of Kdo and components of the CTLH complex across the MZT (Figure 1C). Intriguingly, levels of all components, except for Kaz (one of the catalytic subunits), decreased markedly during early embryogenesis. For instance, Muskelin levels dropped soon after ME31B degradation and were undetectable by the end of the MZT (Figure 1B, C), followed by RanBPM, Hou, and finally Kdo itself. In other words, the subunits of the CTLH complex are additional examples of maternally deposited proteins that are degraded during the MZT, likely contributing to the temporal control of their degradation targets.

The human ortholog of Muskelin is subject to autoubiquitylation (Maitland et al. 2019; Yi et al. 2024), and so we wondered if similar autoregulation occurred in *Drosophila*. Consistent with such a model, in the absence of the E2 Kdo, Muskelin was stabilized (Figure 1C). In embryos depleted of RanBPM, which is predicted to connect Muskelin to the rest of the complex, Muskelin was also stabilized (Figure 1C). Moreover, although Hou was substantially depleted in the absence of RanBPM, the residual Hou present was still degraded, again consistent with a model of autoregulation of the CTLH complex subunits (Figure 1C).

Given that both Muskelin and ME31B levels decrease during the MZT (Figure 1C), we next asked how their interaction changed during the same developmental window. Despite the robust reduction in levels of both ME31B and Muskelin by the end of the MZT, we were able to enrich for both proteins throughout the time course (Figure 1D). In contrast, by the end of the MZT (at 4–5 hours), we were no longer able to detect an interaction between Muskelin and ME31B (Figure 1D, mCherry knockdown). Interestingly, when Kdo was depleted, this interaction was now detectable at the end of the MZT (Figure 1D), perhaps suggesting that the decrease in levels of these proteins might partly lead to the reduced interaction. Thus, not only do the levels of the CTLH complex decrease, but also the remaining complex interacts poorly or not at all with its target ME31B during the MZT. It is tempting to speculate the reduced interaction may in fact trigger the autoregulation of the CTLH complex.

### Muskelin expression is restricted to oogenesis

We next explored the developmental expression of transcripts encoding known components of the CTLH complex (*RanBPM*, *Hou*, *Sou*, *Kaz*, and *Muskelin*), the E2 *Kdo*, and its known targets (*Cup*, *TRAL*, and *ME31B*) using published RNAseq datasets from FlyBase (Figure 2A; Öztürk-Çolak et al. 2024). Although most components showed broad expression across the available tissue atlas, they tended to be highest in the ovary. A notable exception was the component *Muskelin*, whose transcript appears to be highly restricted to the ovary, an observation we confirmed with qRT-PCR (Figure S2A). In fact, when we compared absolute transcript abundances of known CTLH components, *Kdo*, and CTLH targets by Pearson correlation (Figure 2B), *Muskelin* was most similar to the *targets* of the CTLH complex, not other complex members. Similar results were seen when we analyzed published mass spectrometry and RNAseq datasets from FlyBase (Casas-Vila et al. 2017; Graveley et al. 2011; Figures S2B, S2C). Given that Muskelin is required for the degradation of ME31B, Cup, and TRAL (Zavortink et al. 2020), its own restricted developmental expression may limit their degradation.

**Figure 2.**
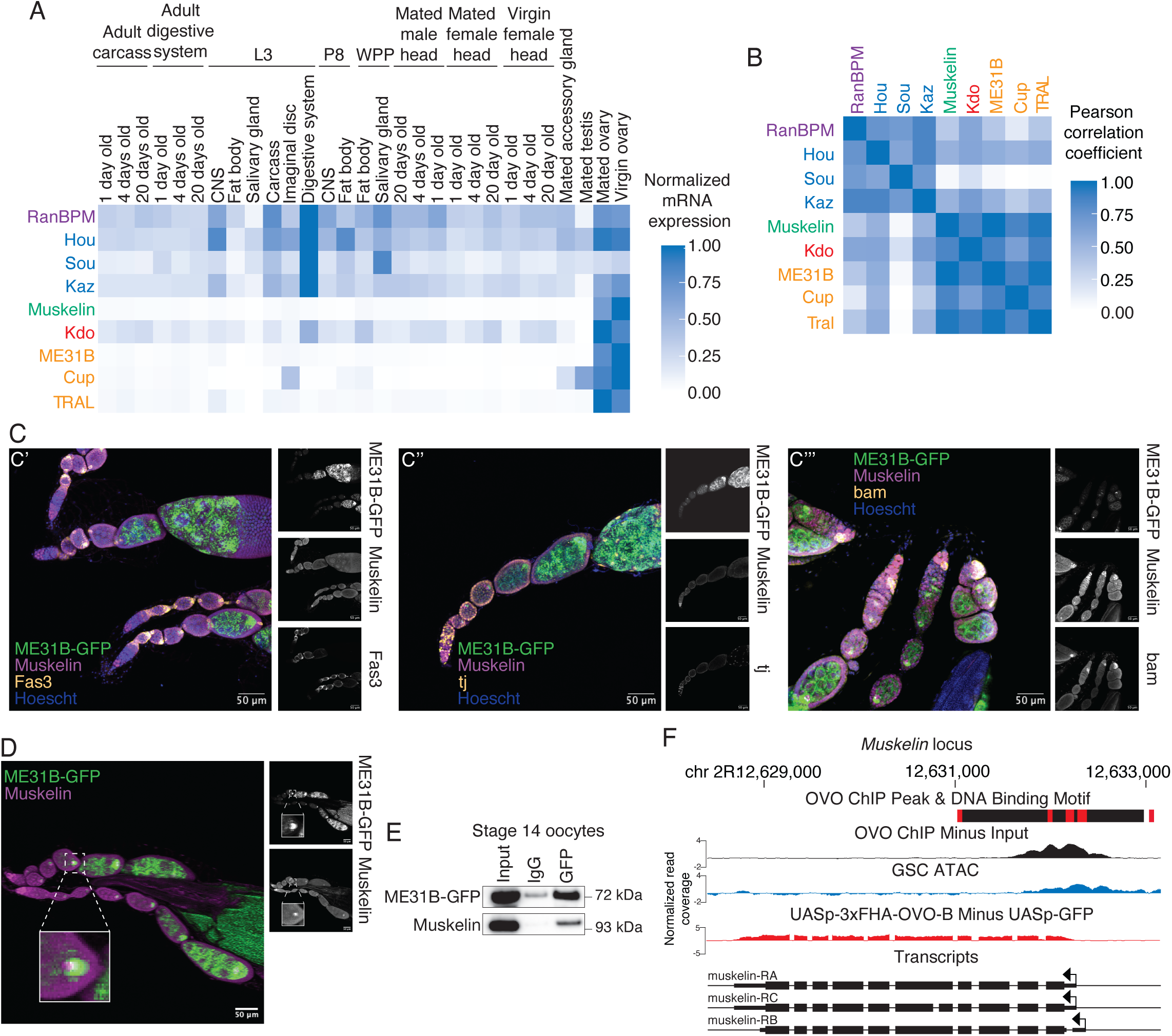
Muskelin expression is restricted to oogenesis. A. mRNA expression of CTLH components and targets is highest in ovarian tissues. Each gene’s expression in the indicated tissue was normalized to its maximum expression. “L3”: third instar larvae; “P8”: pupal stage 8, “WPP”: white prepupa. B. *Muskelin* mRNA expression is most similar to *Kdo* and CTLH complex targets. Pearson correlation coefficients between absolute RPKM values across all tissues from (A) were calculated for each indicated gene pair. C. Muskelin is expressed in a variety of cell types in the ovary. Ovariole co-immunofluorescence was performed for Muskelin (purple) and Fas3, tj, or bam (yellow) in ME31B-GFP (green) expressing ovaries. Images are representative of at least three biological replicates. Muskelin co-localizes with Fas3 (C’) and tj (C’’) in follicle cells and with bam (C’’’) in germline stem cells. Scale bar = 50 µm. D. Muskelin and ME31B-GFP co-localize as puncta in the developing oocyte. Wild-type ovarioles expressing ME31B-GFP (green) were stained for Muskelin (magenta). Germline stem cells are at the left of the image with more mature egg chambers toward the right. ME31B-GFP expression is highest in the developing oocyte where co-localization with Muskelin is strongest. Scale bar = 50 µm. Inset represents a 400% magnification of the indicated oocyte. E. Muskelin and ME31B-GFP interact by immunoprecipitation in stage 14 oocytes. ME31B-GFP was immunoprecipitated from stage 14 oocyte lysate and probed for Muskelin and ME31B-GFP. Representative data from three biological replicates. F. *Muskelin* gene level read coverage tracks for OVO ChIP minus input and GSC ATAC-seq, and *ovo^ΔBP^*/*ovo^ovo-GAL4^*; *UASp-3xFHA-OVO-B* minus *ovo^ΔBP^*/*ovo^ovo-GAL4^*; *UASp-GFP* RNA-seq. Red rectangles and black rectangles represent significant OVO DNA binding motifs and OVO ChIP peaks, respectively. Gene models are represented at bottom. Small rectangles represent untranslated regions, large rectangles represent translated regions. Arrows indicate transcriptional start sites.

These analyses of Muskelin came with two caveats, however. First, whole ovaries are a complex mix of cells. Second, even within the oocyte, complex regulatory networks can translationally repress maternal transcripts until egg activation; indeed, *Kdo* is an example of this phenomenon (Zavortink et al. 2020). In other words, the presence of an mRNA in the oocyte does not necessarily mean the presence of its encoded protein. We next explored the expression of Muskelin using immunofluorescence, probing for the endogenous protein using our newly developed antibody (Figure S2D). In wild-type ovaries, we observed that Muskelin is present in the developing oocyte and is detectable in early germline stem cells and follicle cells (Figure 2C). We used bam as a marker for germline stem cells and Fas3 and traffic jam (tj) as markers for follicle cells (Figure 2C’, 2C’’, 2C’’’; Rust et al. 2020; Li et al. 2003; Newton et al. 2015). The wide expression of Muskelin we observe by immunofluorescence is consistent with single-cell RNAseq data indicating Muskelin expression outside the germline and previous studies showing that it plays roles outside of the developing oocyte to support oogenesis (Li et al. 2022; Kronja et al. 2014).

We next compared the localization of Muskelin with ME31B, using an ME31B-GFP allele expressed at the endogenous locus (Zavortink et al. 2020). We have observed that these proteins interact in the embryo (Figure 1D), and so we were curious whether this interaction is initiated in the oocyte. Indeed, the two proteins colocalized as puncta in the developing oocyte (Figure 2D, Figure S2D). In addition, an interaction between ME31B-GFP and Muskelin was detectable by immunoprecipitation from stage 14 oocytes (Figure 2E). Because Kdo has not yet been produced in the oocyte (Zavortink et al. 2020), the Muskelin-ME31B interaction most likely does not result in ubiquitylation; instead, we propose that the CTLH complex is “poised” in the oocyte prior to egg activation.

Intrigued by our finding that Muskelin is expressed in germline stem cells, we considered potential transcription factors that could activate its expression. Recent work has characterized targets of the master oogenesis transcription factor OVO (Benner et al. 2024). Importantly, the *Muskelin* promoter contains predicted OVO binding sites and OVO ChIP-seq signal (Figure 2F). *Muskelin* transcripts are also observed in ovaries where a hypomorphic *ovo-B* allele has been rescued by OVO overexpression (Figure 2F; Benner et al. 2024). Similarly, *Kdo*, whose transcript levels are also higher in the ovary than other adult tissues (Figure 2A), contains OVO binding motifs, has ChIP-seq signal at the promoter, and is transcriptionally rescued by OVO overexpression in *ovo-B* hypomorphs (Figure S2D). Thus, we conclude that *Muskelin* and *Kdo* are targets of OVO and expressed early in oogenesis, although in the case of *Kdo* its mRNA remains translationally repressed until egg activation (Zavortink et al. 2020; Eichhorn et al. 2016). Together, these results demonstrate that OVO is a key transcription factor for *Muskelin* expression, thus explaining its restriction to the ovary while still being broadly detectable within different cell types in ovarioles.

### Muskelin bridges the interaction between ME31B and the CTLH complex

Given that *Muskelin* expression is developmentally restricted to the ovary, we were curious about its role within the *Drosophila* CTLH complex. Recent work on the human and yeast versions of the CTLH has shown that the WD40 β-propeller-containing components, like Muskelin, often play a structural role in the complex, allowing the generation of a massive, “chelated” E3 ligase (Sherpa et al. 2021; Qiao et al. 2020; Gottemukkala et al. 2024; Yi et al. 2024). At the same time, in humans, WDR26 (which also contains this same domain) has been shown to act as a substrate adaptor for the complex in lieu of the canonical Gid4 subunit (Mohamed et al. 2021; Gross et al. 2024; Maitland et al. 2024). We reasoned that Muskelin might either enable a structural conformation required for interaction with ME31B or, possibly, act as the substrate receptor itself.

We first confirmed, as we have shown previously (Zavortink et al. 2020), that degradation of ME31B-GFP depends upon Muskelin as does the interaction between ME31B and the CTLH complex in 0–1 hour embryos (as monitored by RanBPM; Figures 3A, B). In contrast, different results emerged when RanBPM was depleted (Figure 3C). Although the interaction between Muskelin and other components of the CTLH complex (i.e., Hou and Kaz) was readily detectable in control knock-down embryos, these interactions were no longer detected in the absence of RanBPM. This result is consistent with known structures where RanBPM (and its orthologs) helps connect the WD40 β-propeller subunits to the rest of the complex (van gen Hassend et al. 2023; Sherpa et al. 2022; Lampert et al. 2018). In contrast, Muskelin still interacted with ME31B in the absence of RanBPM. In other words, although Muskelin is required for the interaction between ME31B and the CTLH complex, RanBPM is not required for the interaction between Muskelin and ME31B (Figure 3D).

**Figure 3.**
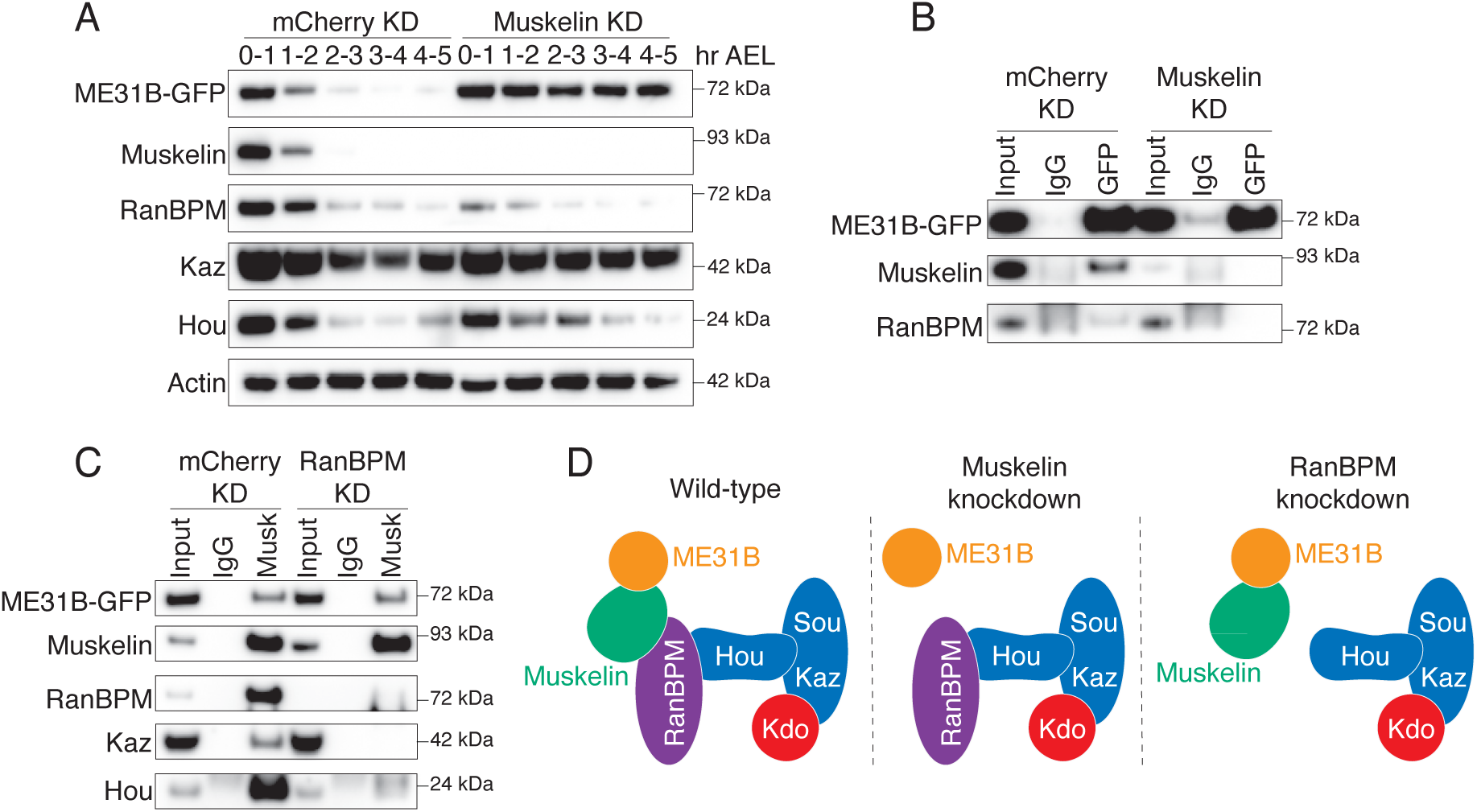
Muskelin bridges the interaction between ME31B and the CTLH complex. A. Muskelin depletion stabilizes ME31B-GFP. ME31B-GFP and CTLH component protein levels were probed in staged lysates from control embryos (mCherry KD) and embryos lacking Muskelin (Muskelin KD). “Hr AEL”: hours after egg laying. B. ME31B-GFP interacts with RanBPM only in the presence of Muskelin. ME31B-GFP was immunoprecipitated from 0-1 hour control (mCherry) or Muskelin knock-down embryo lysate and probed for ME31B-GFP, Muskelin, and RanBPM. C. Muskelin and ME31B-GFP interact independently of the CTLH complex. Muskelin was immunoprecipitated from lysates from staged 0-1 hour control (mCherry) or RanBPM knock-down embryos, and samples were blotted for CTLH complex components and ME31B-GFP. All western blots are representative of three biological replicates. D. Muskelin bridges the interaction between ME31B and the CTLH complex via its interaction with RanBPM, and RanBPM is not required for the interaction between Muskelin and ME31B.

### Muskelin is the substrate recognition adaptor of the embryonic CTLH complex

Given the lack of a known Gid4 substrate receptor homolog in flies, we explored the possibility of an unknown component of the embryonic CTLH complex that acts in this role and employed a series of complementary immunoprecipitation/mass spectrometry experiments and analysis (Figure 4A–C; Supp Tables 1-3). First, we used Hou as bait to define the embryonic complex. We confirmed that Hou immunoprecipitations from 0–1 hour embryo lysate contained RanBPM, Muskelin, Kaz, and ME31B by western blotting, indicating that our antibody was able to pull-down Hou in the context of the full complex (Figure S3A). Next, we performed mass spectrometry on immunoprecipitated proteins (Table S1). As expected, relative to the IgG control, known targets (ME31B, Cup, and TRAL) were significantly enriched in the pull-downs as were previously identified components of the CTLH complex (Figure 4A). YPEL5 (Yippee in flies) was identified in the Hou pulldown, which is a known homolog of a known human CTLH component that acts to inhibit substrate ubiquitylation (Gottemukkala et al. 2024). However, this protein was not required for degradation of ME31B nor did its absence appear to substantially enhance ME31B degradation (Figure S3B). Although there were several other significantly enriched proteins in the Hou pulldown, none were homologs of known substrate adaptors nor had an obvious role or connection with the CTLH complex, protein degradation, or ME31B, based on annotations in FlyBase (Table S1).

**Figure 4.**
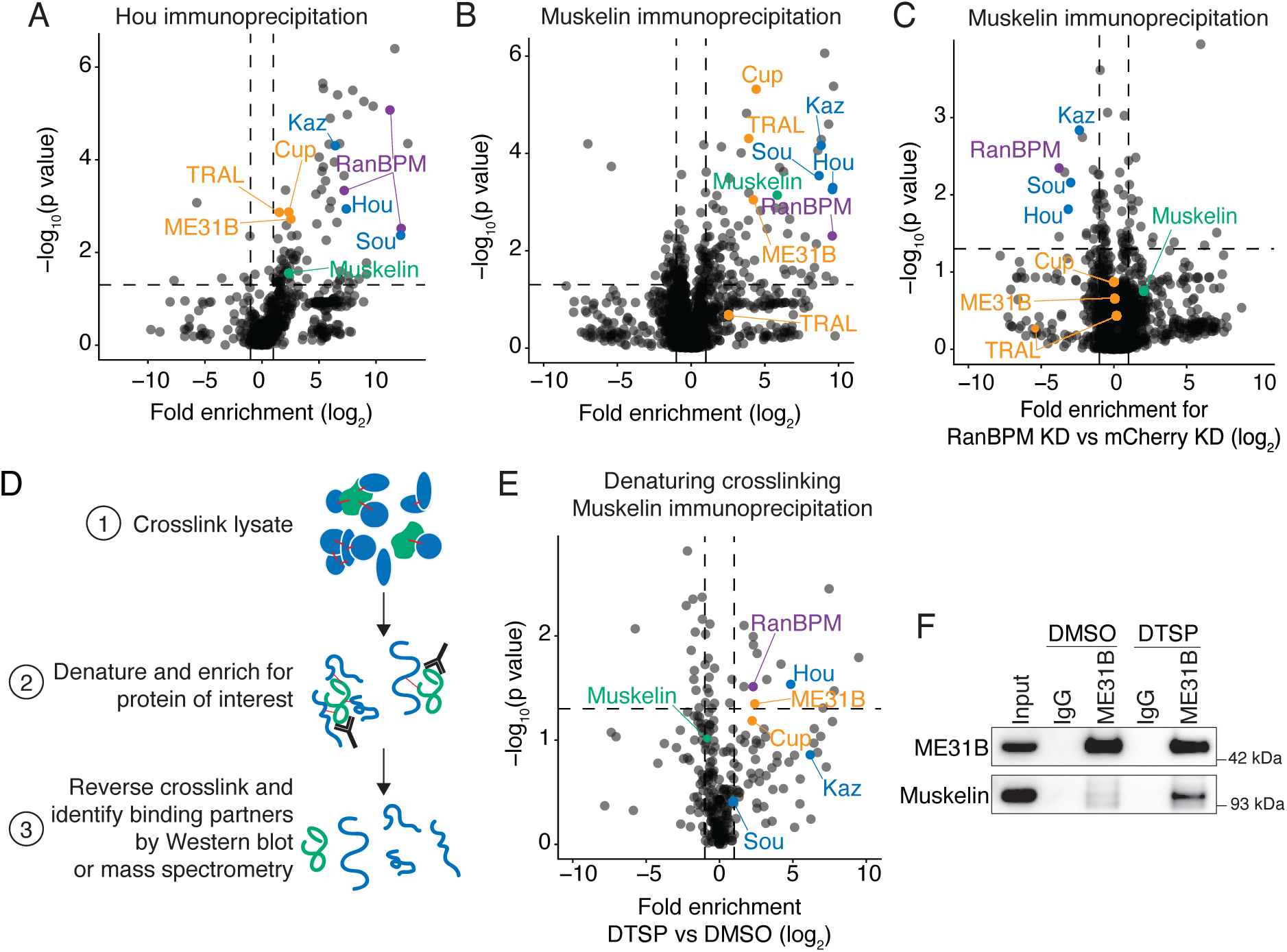
Muskelin is the substrate recognition adaptor of the embryonic CTLH complex. A. Hou immunoprecipitation enriches CTLH complex components and targets. Hou was immunoprecipitated from 0-1 hour embryo lysate and binding partners were identified by mass spectrometry. Fold enrichment over control immunoprecipitation was calculated and plotted against adjusted p-value across three replicates. B. Muskelin immunoprecipitation enriches CTLH complex components and targets. Muskelin was immunoprecipitated from lysates from control 0-1 hour embryos and binding partners were identified by mass spectrometry. Fold enrichment over control immunoprecipitation was calculated and plotted against adjusted p-value across three replicates. C. RanBPM depletion disrupts the interaction between Muskelin and the CTLH complex but not its targets. Muskelin was immunoprecipitated from 0-1 hour control embryo lysate or embryo lysate lacking RanBPM, and binding partners were identified by mass spectrometry. The fold enrichment of each protein in RanBPM knock-down compared to control knock-down samples was calculated and plotted against adjusted p-value across three replicates. D. Denaturing crosslinking immunoprecipitation workflow. Lysates are crosslinked with DTSP, denatured and immunoprecipitated, then the crosslink is reversed, and binding partners are identified. E. Denaturing crosslinking Muskelin immunoprecipitation identifies CTLH components and targets as close-proximity binding partners in 0-1 hour *w1118* embryos by mass spectrometry. Fold enrichment of each protein from a Muskelin immunoprecipitation in DTSP-treated (crosslinked) lysate vs DMSO-treated (uncrosslinked) lysate was calculated and plotted against adjusted p-value across three replicates. F. The ME31B-Muskelin interaction is detectable only in crosslinked samples from a denaturing crosslinking ME31B immunoprecipitation. ME31B was immunoprecipitated from 0-1 hour *w1118* embryo lysate under denaturing conditions after crosslinking with DTSP or DMSO as a control. Lysates were then probed for ME31B and Muskelin. All western blots are representative of three biological replicates.

Second, we cross-referenced all significantly enriched proteins from published Muskelin-GFP and ME31B-GFP immunoprecipitation mass spectrometry datasets (Cao et al. 2020; Zavortink et al. 2020). Given that a putative, unknown substrate adaptor would likely interact with both Muskelin and ME31B, we reasoned it should be present in both datasets (which robustly detected the ME31B-Muskelin interaction). Once again, no homologs of known substrate adaptors were present in the list of high-confidence enriched targets. We further cross-referenced the significantly enriched proteins between the same Muskelin-GFP immunoprecipitation dataset with our Hou immunoprecipitation mass spectrometry dataset (Figure S3C). None of the proteins present in both datasets were obvious homologues of CTLH components or had a role in protein degradation (Table S2).

Finally, we immunoprecipitated Muskelin from 0–1 hour embryo lysate that lacked RanBPM and performed mass spectrometry (Table S3). Consistent with our previous results (Figure 2C), we were able to see significant enrichment of ME31B, Cup, and TRAL relative to control pulldowns (Figure 4B). Also consistent with previous results, in the Muskelin immunoprecipitation from embryos lacking RanBPM, the interaction with the rest of the CTLH complex was lost despite maintaining an interaction with ME31B, Cup, and TRAL (Figure 4C). However, as before, no potential alternative substrate receptor was enriched.

Given these findings, we next considered the intriguing idea that Muskelin acts as the substrate receptor for the CTLH complex and explored the potential for a direct interaction between it and ME31B. To do so, we optimized denaturing crosslinking immunoprecipitations from embryo lysates (Figure 4D). This protocol uses Dithiobis(succinimidyl propionate) (DTSP), a bivalent covalent protein crosslinker, to reversibly link all lysine residues within 12 Angstroms of one another (Akaki et al. 2022). We crosslinked native early embryo lysate and then enriched for proteins crosslinked to Muskelin by performing immunoprecipitations under denaturing conditions with stringent washes. After reversing the crosslinks, immunoprecipitated proteins were identified by mass spectrometry in three biological replicates. RanBPM was significantly enriched, as was Hou (Figure 4E, Table S4). Strikingly, ME31B was also significantly enriched compared to the non-crosslinking control (Figure 4E, Table S4). Cup was enriched but did not meet our significance cut-off of corrected *p-*value < 0.05. When we performed the reverse experiment, now immunoprecipitating ME31B under denaturing conditions, we were able to detect Muskelin only in the presence of DTSP (Figures 4F, S3D; Table S5). Thus, we conclude that Muskelin directly interacts with ME31B and serves as the substrate receptor for the embryonic CTLH complex in *Drosophila*.

### The main targets of the embryonic CTLH complex are ME31B, Cup, and TRAL

To investigate what other proteins were targeted by Muskelin and the CTLH complex, we performed total protein mass spectrometry on embryos from control, RanBPM-depleted, or Muskelin-depleted embryos in the early and late MZT (Table S6). We would expect that in wild-type embryos, many proteins will be degraded regardless of their dependence on the CTLH complex (Figure 5A). However, in Muskelin knockdown embryos, we would expect that only specific targets of Muskelin to be stabilized, while Muskelin-independent CTLH targets still degraded. Finally, in RanBPM knockdown embryos, we would expect that all CTLH targets to be stabilized, including Muskelin-dependent targets.

**Figure 5.**
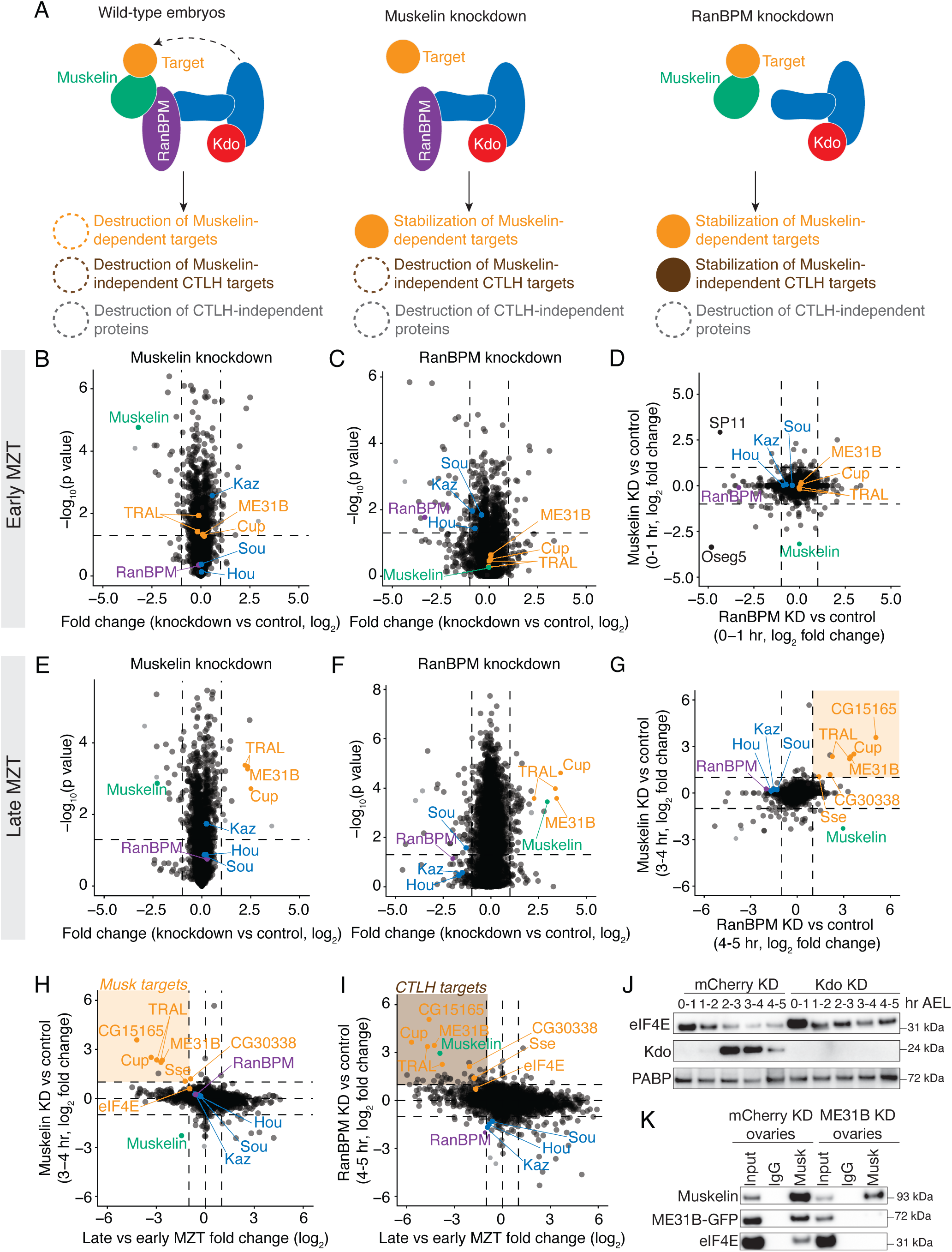
The main targets of the embryonic CTLH complex are ME31B, Cup, and TRAL. A. Muskelin regulates a distinct subset of the CTLH-dependent proteome. In wild-type embryos, many types of proteins are degraded that are both dependent on and independent of CTLH activity. In Muskelin knock-down embryos, only proteins that are specifically targeted by Muskelin are stabilized, while other CTLH targets are still degraded. In RanBPM knock-down embryos, all CTLH target proteins are stabilized, including those targeted by Muskelin. B. Lack of Muskelin in the early embryo has minor effects on the proteome. The fold changes of protein levels in 0-1 hour Muskelin knock-down embryos compared to control knock-down embryos were plotted against adjusted p-value. All points represent the average fold change of a specific gene across three replicates. C. Lack of RanBPM in the early embryo has minor effects on the proteome. Whole-proteome fold changes of 0-1 hour RanBPM knockdown embryos were compared to 0-1 hour control embryos and plotted against adjusted p-value across three replicates, otherwise as in (A.) D. There are few proteomic differences between early embryos lacking RanBPM and those lacking Muskelin. Whole-proteome fold changes of 0-1 hour Muskelin knockdown embryos were compared to those of 0-1 hour RanBPM knockdown embryos across three replicates. E. ME31B, Cup, and TRAL are significantly stabilized in late-MZT embryos lacking Muskelin. Levels of other CTLH components do not change substantially. Whole-proteome fold changes of 3-4 hour Muskelin knockdown embryos were compared to 3-4 hour control embryos and plotted against adjusted p-value across three replicates. F. Muskelin, ME31B, Cup, and TRAL are significantly stabilized in late-MZT embryos lacking RanBPM. Whole-proteome fold changes of 4-5 hour RanBPM knockdown embryos were compared to 4-5 hour control embryos and plotted against adjusted p-value across three replicates. G. A small subset of the proteome is stabilized in the absence of RanBPM or Muskelin (orange box). Whole-proteome fold changes of 3-4 hour Muskelin knockdown embryos were compared to 4-5 hour RanBPM knockdown embryos across three replicates. H. Few proteins that are stabilized in embryos lacking Muskelin are also degraded in wild-type embryos during the MZT (orange box). Proteins stabilized in 3-4 hour Muskelin knockdown embryos (see Figure 5D) were plotted against proteins degraded at the end of the MZT in control embryos. I. Few proteins that are stabilized in embryos lacking RanBPM are also degraded in wild-type embryos during the MZT (brown box). Proteins stabilized in 4-5 hour RanBPM knockdown embryos (see Figure 5E) were plotted against proteins degraded at the end of the MZT in control embryos. J. eIF4E is mildly stabilized in embryos depleted of Kdo. eIF4E, Kdo and PABP (as a loading control) protein levels were probed in staged lysates from control embryos and embryos lacking Kdo. “Hr AEL”: hours after egg laying. K. The interaction between Muskelin and eIF4E depends on ME31B. Muskelin was immunoprecipitated from control ovary lysate or ovary lysate lacking ME31B and probed for Muskelin, ME31B-GFP, and eIF4E. All western blots are representative of three biological replicates.

Early in the MZT (at 0–1 hour), there were very few changes between Muskelin knockdown and control (Figure 5B), and only 15 proteins changed by at least 2-fold (with a corrected *p-*value < 0.05). Consistent with a low level of Kdo at this point in embryogenesis, none of the known targets of the CTLH complex (ME31B, Cup, and TRAL) were significantly different. Similar results were observed in embryos depleted of RanBPM at 0-1 hour (Figure 5C, Table S6). Interestingly, several extracellular proteins (Oseg5 and SP11) were depleted in embryos lacking either Muskelin or RanBPM at this timepoint, perhaps reflecting defects that have been previously observed when Muskelin is depleted from the ovary (Kronja et al. 2014; Figure 5D).

We next examined how loss of Muskelin or RanBPM affected the proteome at the end of the MZT (3-4 and 4-5 hours after egg laying, respectively; Figures 5E–G). Consistent with our analysis of the autoregulation by the CTLH complex, RanBPM, Kaz, Sou, and Hou were not stabilized during the MZT in the absence of Muskelin (Figure 5E), but Muskelin was significantly stabilized in the absence of RanBPM (Figure 5F). Moreover, and again as expected, Cup, ME31B, and TRAL were robustly stabilized at these time points in the absence of either Muskelin or RanBPM relative to control knockdowns (Figures 5E, F). Surprisingly, however, only 12 and 28 other proteins were upregulated in the Muskelin and RanBPM knockdown embryos, respectively (Figures 5E, F). Of these, only eight were common to both datasets, three of which were Cup, ME31B, and TRAL (Figure 5G). Intrigued by the possibility that these other proteins might represent targets of the CTLH complex, we compared how levels of these proteins changed during the MZT with how they change upon loss of the E3 ligase component (Figures 5H, I). Three of them (CG15165, CG30338, and Sse) were both destroyed during the MZT and stabilized upon the loss of either Muskelin or RanBPM. Nonetheless, of these putative additional substrates, none were enriched in any Muskelin pulldown (Table S3, Table S4), indicating that Muskelin does not serve as the substrate adaptor for these specific proteins’ clearance. Along these lines, it is notable that there appear to be more proteins that are degraded in wild-type embryos which are stabilized in embryos lacking RanBPM than are stabilized in embryos lacking Muskelin, perhaps indicating Muskelin-independent CTLH targets.

We noticed that eIF4E, which interacts with ME31B and its binding partners, was also slightly stabilized in the absence of RanBPM or Muskelin (Figure 5H, I), although it failed to reach our cut-off thresholds. To investigate this observation, we probed levels of eIF4E across the MZT in control or Kdo knock-down embryos. As has been observed previously (Cao et al. 2020), eIF4E levels modestly decreased during the MZT, but were stabilized in embryos depleted of Kdo (Figure 5J). However, the effect was much weaker than that of ME31B, Cup, and TRAL (Figures 5H, I), leading us to wonder if it was directly recognized by the CTLH complex. To explore this idea, we immunoprecipitated Muskelin from control ovary lysate and lysate lacking ME31B. Although we detected an interaction between eIF4E and Muskelin in control ovaries, this interaction was not detectable upon loss of ME31B (Figure 5K). This result suggests that the CTLH complex may not directly recognize eIF4E and that its degradation may instead be mediated by its proximity to ME31B. Taken together, these data indicate that the embryonic CTLH complex via its adaptor Muskelin regulates a very small number of proteins, and ME31B, Cup, and TRAL appear to be its major targets.

## DISCUSSION

We have found multiple transcriptional and post-translational regulatory mechanisms restrict the degradation of ME31B to early embryogenesis (Figure 6). Previously, we had found that the corresponding E2 conjugating enzyme for the CTLH complex, Kdo, was translationally controlled such that it was only produced upon egg activation (Zavortink et al. 2020). Indeed, we have now shown that the CTLH complex is able to bind its target ME31B in the mature oocyte via Muskelin, but this interaction cannot result in ubiquitylation until Kdo is produced at the onset of the MZT. In other words, in the oocyte, the CTLH complex appears poised to degrade its targets once translation of *Kdo* is triggered.

**Figure 6.**
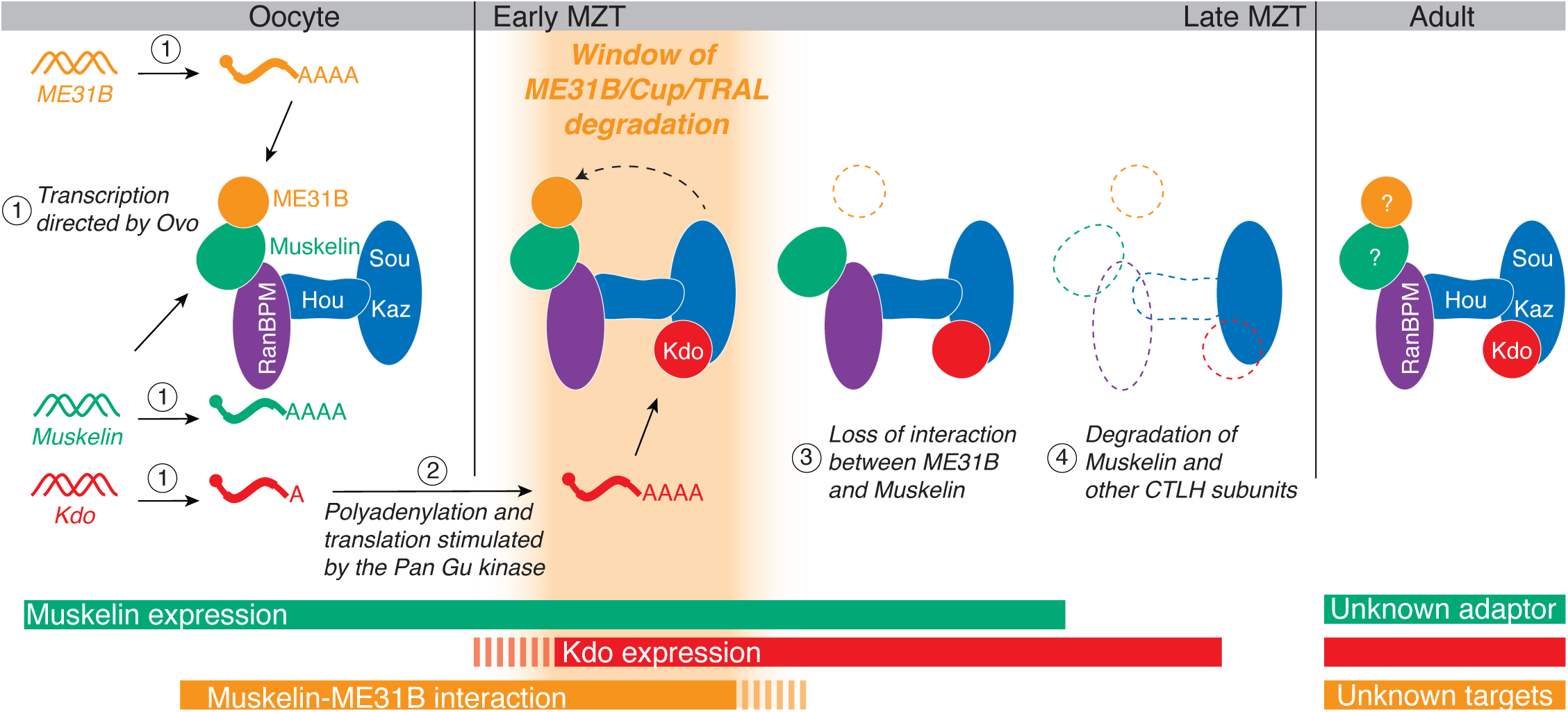
Model: At least four mechanisms restrict the degradation of ME31B and its binding partners Cup and TRAL to the MZT. First, in oocytes, transcription of *Muskelin, ME31B, and Kdo* is orchestrated by the oocyte-specific transcription factor OVO (1). *Kdo* remains translationally repressed until egg activation by activity of the Pan Gu kinase (2), but Muskelin and ME31B are translated in the oocyte. Muskelin bridges the interaction between ME31B and the CTLH complex in the developing oocyte, but because Kdo is not yet translated, the complex is “poised” and inactive. At egg activation and the beginning of the MZT, *Kdo* mRNA is translated, and its increasing protein levels activate the CTLH complex, enabling the ubiquitylation that leads to ME31B/Cup/TRAL degradation (depicted as only ME31B for clarity). At the end of the MZT, Muskelin and ME31B no longer interact (3). Finally, Muskelin, Kdo, much of the CTLH complex, and CTLH targets are degraded, effectively stopping additional degradation (4). After the MZT in the adult fly, CTLH components and Kdo are still present, but the substrate adaptor and targets in this context remain unknown. These overlapping mechanisms restrict the window of ME31B degradation to a small timeframe during the MZT.

Importantly, we found that Muskelin acts as the substrate receptor for the embryonic CTLH complex. This role for Muskelin contrasts with our current understanding of the human complex, where it appears to primarily mediate multimerization of the complex rather than direct substrate recognition (van gen Hassend et al. 2023). Like many substrate adaptors, Muskelin itself is tightly regulated, and its regulation appears to be a major mechanism restricting the degradation of ME31B to the MZT. The expression of *Muskelin* transcripts is remarkably specific to the ovary. Our analysis points to its expression in germline stem cells, follicle cells, and the growing oocyte itself. A large part of its expression is due to OVO, a critical transcription factor in the female germline. Analysis of available datasets (Benner et al. 2024) indicate that the *Muskelin* promoter contains OVO binding motifs and is indeed bound by OVO by ChIP-seq. It is interesting to note that OVO also controls *ME31B*, *Cup*, *TRAL*, and *Kdo* mRNA expression (Benner et al. 2024), demonstrating another common regulatory mechanism between the CTLH complex and its targets.

The CTLH complex is also post-translationally controlled. We observed that in the oocyte and early MZT, the interaction between Muskelin and ME31B is robust, but is no longer detectable by the end of the MZT. Similarly, the CTLH complex itself is degraded at the end of the MZT, and levels of nearly every member of the CTLH complex, as well as Kdo, decrease at this time in development. Given that protein degradation relies upon the E2 Kdo and that human Muskelin is subject to autoregulation (Maitland et al. 2019), we propose that the embryonic *Drosophila* CTLH complex is similarly controlled and that it is triggered by the loss of interaction with its targets. In support of such a mechanism, the degradation of Muskelin is dependent on RanBPM. We also found evidence that RanBPM and Hou are co-regulated because the depletion of RanBPM led to a loss of Hou as well by western blotting (Figure 1C). There is precedent for this finding: other work has shown that knockdown of the human ortholog of Hou (TWA1) largely destabilizes the entire CTLH complex (Lampert et al. 2018) and that TWA1 must be present in order to purify murine RanBPM (RanBP9; van gen Hassend et al. 2023).

Another surprise from our study is that, despite this complex regulatory network, the embryonic Muskelin-CTLH complex appears to have a limited number of targets, and our analysis suggests that the most important targets are Cup, ME31B, and TRAL. These three proteins are critical for oogenesis and help mediate the post-transcriptional repression of thousands of transcripts in the oocyte and embryo (Wang et al. 2017). Given that the role of the poly(A) tail and decapping change during the MZT (Wang et al. 2017), it is tempting to speculate that the degradation of ME31B-Cup-TRAL (which bind the mRNA cap binding protein eIF4E) may be mechanistically linked to resetting the post-transcriptional landscape. Nonetheless, understanding the impact of their degradation on both maternal and zygotic mRNAs remains an important and unresolved issue.

Even though this work has expanded our understanding of maternal protein degradation during the MZT and the CTLH E3 ligase, it has also revealed new compelling questions. For instance, the degron recognized by Muskelin is unknown, as is whether Muskelin recognizes the ME31B-TRAL-Cup complex as a whole. That eIF4E is degraded seemingly because of its proximity to ME31B suggests that perhaps the entire complex is recognized. If so, understanding the higher order structure of the embryonic CTLH complex becomes more critical – how does the E3 ligase accommodate such a large protein complex, and, as a related issue, is this the mRNA-bound form? Another unanswered question is the identity or presence of other active substrate adaptors for the CTLH complex during and outside the MZT. One tempting possibility for an additional substrate adaptor is *CG7611*, the *Drosophila* ortholog of human

*WDR26*. This subunit acts as the substrate adaptor in specific contexts in humans, but is not required for the degradation of ME31B and its binding partners (Gross et al. 2024; Gottemukkala et al. 2024; Zavortink et al. 2020). With recent work highlighting the dual roles of Muskelin and WDR26 (Onea et al. 2022; Maitland et al. 2024), our work provides a fascinating perspective on the evolutionary plasticity between structural components and substrate adaptors in the CTLH complex. Our results motivate further investigations into the make-up and organization of this otherwise well-conserved E3 ligase in multiple species.

## MATERIALS AND METHODS

### Drosophila fly stocks

Fly stocks were maintained at 22°C with 50% humidity. OVO fly lines were generated and maintained according to Benner et al. 2024 and maintained by the Oliver lab. All knockdowns were generated from commercially available UAS-RNAi lines crossed to a ME31B-GFP;7062 Gal4 driver (Zavortink et al. 2020). RNAi lines: mCherry (BDSC 35785), Kdo (BDSC 51410), RanBPM (BDSC 61172), Muskelin (BDSC 51405).

### S2 cells

Schnieder’s S2 cells (ThermoFisher Scientific) were grown at 28°C and maintained in media containing Express Five™ SFM (ThermoFisher Scientific) with 10% L-glutamine.

### S2 cell RNAi

cDNA was generated by amplifying coding region of interest (obtained as plasmids from DGRC) with bidirectional T7 promoters (Table S7). cDNA was transcribed using HiScribe T7 High Yield RNA Synthesis Kit (NEB) according to manufacturer’s directions. RNA was annealed slowly overnight, and dsRNA was transfected using Effectene (Qiagen) according to manufacturer’s instructions. Cells were harvested and lysed in Lysis Buffer A after 72 hours.

### Ovary and ovariole isolation

Ovaries and ovarioles were harvested from 1-3 day-old virgin *Drosophila* according to (Merkle 2023).

### Embryo and ovary lysate preparation

Embryos were collected at various time points post egg-laying on standard yeasted apple juice-agar plates at 25°C. Embryos were dechorionated in bleach, washed in PBS, homogenized in lysis buffer A (150 mM KCl, 20 mM HEPES-KOH pH 7.4, 1 mM MgCl_2_, 1 mM DTT, complete mini EDTA-free protease inhibitors), and were clarified at 14,000 rpm, 4°C for 5 min. Lysates were diluted to 1 mg/mL and stored at −80°C. Ovaries were homogenized in lysis buffer A and clarified as above.

### Antibody production

Antibodies from Boster Bio were produced using known protein sequences from UniProt, accessed via FlyBase.

### RT-qPCR

RNA was extracted using TRIzol (ThermoFisher) according to manufacturer’s instructions. cDNA was generated using SuperScript III (ThermoFisher) according to manufacturer’s instructions and amplified with iTaq SYBR reagents (Bio-RAD) using gene-specific primers (Table S7) and normalized to *Act5C*.

### Immunoprecipitations

Lysates were incubated with GFP (Roche), Muskelin (Boster), Houki (Boster), or IgG (Abcam) antibodies and rotated for 1 hour at 4°C. EZView protein G affinity beads (Sigma) were washed with lysis buffer A 3x times. The lysate-antibody mixtures were added to 25 uL of washed protein G beads and rotated for 1 hour at 4°C. Beads were washed 3x with lysis buffer A and transferred to new tubes. Reducing agent (NuPAGE) and sample buffer (NuPAGE) were added to the beads prior to western blots.

### Denaturing crosslinking immunoprecipitations

Lysates were incubated with 50 mM DTSP (Thermo Fisher) or DMSO (Thermo Fisher) for 3 hours at 4°C. The reaction was quenched by adding Tris-HCl pH 7.5 to 20 mM. An equal volume 2X denaturing lysis buffer (2X PBS, 2% Triton X-100, 2% sodium deoxycholate, 2% SDS) was added followed by immunoprecipitation with ME31B (Boster), Muskelin (Boster), or IgG (Abcam) for 1 hour at 4°C. EZView protein G affinity beads (Sigma) were washed with lysis buffer A 3x. The lysate-antibody mixtures were added to 25 uL of washed protein G beads and rotated for 1 hour at 4°C. Beads were washed 3x with denaturing wash buffer (50 mM Tris pH 8, 1M NaCl, 1% NP-40, 0.05% sodium deoxycholate, 0.1% SDS) and transferred to new tubes. Samples were reduced with DTT to reverse crosslink prior to western blotting or mass spectrometry analysis.

### Western blotting

Samples were run on standard Bis-Tris 4-12% gels (Invitrogen) and transferred to PVDF membrane prior to western blotting. Blots were blocked in 5% milk in PBST prior to incubation with primary antibodies: Rabbit anti-Kdo (Pacific Immunology) was used at 1:1000. Rabbit anti-Muskelin (Boster) was used at 1:5000. Rabbit anti-Katazuke (Boster) was used at 1:5000. Rabbit anti-Houki (Boster) was used at 1:500. Rabbit anti-ME31B (Boster) was used at 1:5000. Rabbit anti-RanBPM (Boster) was used at 1:5000. Mouse anti-GFP (Roche) was used at 1:1000. Rabbit anti-eIF4E (Boster) was used at 1:10000. Rabbit anti-PABP (Boster) was used at 1:10,000.

### Immunostaining and microscopy

Ovaries and ovarioles were immunostained according to Merkle et al 2023. Briefly, whole ovaries were fixed in 4% paraformaldehyde in PBST for 15 minutes at room temperature. Following fixation, they were blocked in 5% BSA in PBST at 4°C overnight. Ovaries were washed with PBST, then primary antibody was added for 4C overnight (rabbit anti-Muskelin 1:500 [Boster], mouse anti-Fas3 1:100 [DHSB], mouse anti-bam 1:250 [DHSB], guinea pig anti-Tj 1:2000 [gift from Oliver lab]). Following a second wash, fluorescent secondaries were added overnight at 4°C (goat-anti rabbit AlexaFluor 647 1:500, goat anti-mouse AlexaFluor 568 1:500, goat anti-guinea pig AlexaFluor 594 1:500; all ThermoFisher). Ovarioles were hand-dissected from whole fixed ovaries prior to imaging.

Ovarioles were then prepared for imaging by embedding in 0.2% agar in PBS on glass bottom imaging dishes (Ibidi). Samples were imaged with a 10X objective on a Stellaris 8 line scanning confocal microscope. Image processing was performed in FIJI.

### Co-IP Mass Spectrometry

#### On-bead digestion

Beads were washed 3 times with 50 mM HEPES pH 8.7 and suspended in 50 mM HEPES pH 8.7. Disulfide bonds within the proteins were reduced by the addition of tris (2-carboxyethyl) phosphine (TCEP) to a concentration of 5 mM with mixing at room temperature for 15 min. The reduced proteins were alkylated by the addition of 2-chloroacetamide to a concentration of 10 mM with mixing in the dark at room temperature for 30 min. Excess 2-chloroacetamide was quenched by the addition of dithiothreitol to a concentration of 10 mM and mixing at room temperature for 30 min. Samples were digested by the addition of 500 ng of rLys-C with mixing at 37 ℃ for 4 hours followed by the addition of 500 ng of trypsin protease and mixing overnight at 37 ℃. Samples were filtered over 2 μm spin cartridges and dried in a SpeedVac. Samples were desalted over TopTips (LC Packings), eluted with 15 mM ammonium formate pH 2.8, and dried in a SpeedVac. Samples were resuspended in 20 µL 2% acetonitrile/0.1% formic acid in preparation for LC/MS analysis.

#### TMT multiplexed sample preparation

A streamlined TMT protocol was followed for multiplexed quantitative proteomics experiments (Navarrete-Perea et al. 2018). In short, initial solutions containing 100 μg of protein were in 100 μL of a buffer containing 20 mM HEPES, 100 mM KCl, 0.1 mM EDTA, 0.4% IGEPAL, 10% glycerol, 1 mM dithiothreitol, and protease inhibitor. Disulfide bonds were reduced by the addition of TCEP to a concentration of 5 mM with mixing at room temperature for 15 min. Protein alkylation was carried out by the addition of 2-chloroacetamide to a concentration of 10 mM with mixing in the dark at room temperature for 30 min. Samples were then subjected to protein precipitation using methanol/chloroform. Precipitated protein pellets were resuspended in 70 μL of 50 mM HEPES pH 8.7 and labeled with TMTpro reagent (ThermoFisher Scientific) following the manufacturer’s instructions. Upon completion of labeling, 2 μL from each sample was combined to use as a label check. The resulting pool was briefly placed in a SpeedVac to remove acetonitrile, desalted on an Ultra-micro SpinColumn (Harvard Biosciences), and the elution was taken to dryness. The pool was resuspended in 2% acetonitrile/0.1% formic acid and analyzed on an Orbitrap Fusion to measure the extent of labeling. With the labeling measured to be greater than 97% complete, each labeling reaction was quenched by the addition of hydroxylamine to a final concentration of 0.5%. Samples were combined by adding equal amounts of protein from each sample to the final pool based on total reporter ion intensity measured in the labeling check. The resulting pool was placed in a SpeedVac to remove acetonitrile followed by desalting on a SepPak 200 mg C18 desalting cartridge (Waters) and taken to dryness. The combined TMT sample was resuspended in 100 µL of 10 mM ammonium bicarbonate pH 8. The material was loaded on to a Zorbax 2.1 mm x 150 mm (5 µm particle size) Extend-C18 column (Agilent) for basic reverse-phase fractionation. Ninety-six fractions were collected and combined into 24 pools by concatenation. The pools were subsequently taken to near-dryness by vacuum centrifugation and brought up in 20 µL 2% acetonitrile/0.1% formic acid in preparation for LC/MS analysis.

### Mass spectrometry analysis

Samples from co-immunoprecipitation experiments were analyzed by LC/ESI MS/MS with a Thermo Scientific Easy1200 nLC (Thermo Scientific) coupled to an Orbitrap Fusion, Orbitrap Eclipse with FAIMS Pro (Field Asymmetric Ion Mobility Spectrometry), or Orbitrap Ascend with FAIMS Pro Duo (Orbitraps are manufactured by Thermo Scientific) mass spectrometer. In-line desalting was accomplished using a reversed-phase trap column (100 μm × 20 mm) packed with Magic C_18_AQ (5-μm 200 Å resin; Michrom Bioresources, Auburn, CA) followed by peptide separations on a reversed-phase column (75 μm × 270 mm) packed with ReproSil-Pur C_18_AQ (3-μm, 120 Å resin; Dr. Maisch, Baden-Würtemburg, Germany) directly mounted on the electrospray ion source. A 90-minute gradient from 8% to 30% B (80% acetonitrile/0.1% formic acid) at a flow rate of 300 nL/min was used for chromatographic separations. LC/MS/MS data were collected using data dependent acquisition methods with the MS survey scans detected in the Orbitrap. MS/MS spectra were detected in the linear ion trap using HCD activation. Selected ions were dynamically excluded for 60 seconds after a repeat count of 1. Data analysis was performed using Proteome Discoverer 2.5 (Thermo Scientific). The data were searched against a Uniprot Drosophila melanogaster (UP000000803 downloaded 3-07-21) protein database that included common contaminants (Mellacheruvu et al. 2013). Searches were performed with settings for the proteolytic enzyme trypsin. Maximum missed cleavages were set to 2. The precursor ion tolerance was set to 10 ppm and the fragment ion tolerance was set to 0.6 Da. Dynamic peptide modifications included oxidation (+15.995 Da on M). Dynamic modifications on the protein terminus included acetyl (+42.011 Da on N-terminus), Met-loss (−131.040 Da on M) and Met-loss+Acetyl (−89.030 Da on M) and static modification of carbamidomethyl (+57.021 on C). Minora was used for peak abundance and retention time alignment. Sequest HT was used for database searching. All search results were run through Percolator for scoring and the false discovery rate was set to 1% at the peptide level. Raw quantitative results were transformed to log_2_ scale and normalized to the median value across samples. Missing data were imputed with half of the global minimum value. P-values for pairwise comparisons were calculated by t-test.

Multiplexed quantitative MS data were collected on a ThermoScientific Easy1200 nLC coupled to an Orbitrap Eclipse with FAIMS Pro (ThermoScientific) mass spectrometer. In-line desalting was accomplished using a reversed-phase trap column (100 μm × 20 mm) packed with Magic C_18_AQ (5-μm, 200 Å resin; Michrom Bioresources) followed by peptide separations on a reversed-phase column (75 μm × 270 mm) packed with ReproSil-Pur C_18_AQ (3-μm, 120 Å resin; Dr. Maisch, Baden-Würtemburg, Germany) directly mounted on the electrospray ion source. A 180-minute gradient from 4% to 44% B (80% acetonitrile/ 0.1% formic acid) at a flow rate of 300 nL/min was used for chromatographic separations. The FAIMS Pro source used varied compensation voltages of −40 V, −60 V, −80 V while the Orbitrap Eclipse instrument was operated in the data-dependent mode. MS survey scans were collected in the Orbitrap (resolution 120,000) with a 3 sec cycle time and MS/MS spectra acquisition were detected in the linear ion trap with CID activation using turbo speed scan. Selected ions were dynamically excluded for 60 seconds after a repeat count of 1. Following MS2 acquisition, real time searching (RTS) was employed, and spectra were searched against a Drosophila melanogaster (UP000000803) protein database using the real-time search algorithm COMET to identify suitable MS/MS ions for synchronous precursor selection (SPS) MS3 quantitation. SPS-MS3 was collected on the top 10 most intense ions detected in MS2 and subjected to higher energy collision-induced dissociation (HCD) for fragmentation and subsequent analysis in the Orbitrap (resolution 50,000). Data analysis was performed using Proteome Discoverer 2.5 (Thermo Scientific). The data were searched against a Drosophila melanogaster (UP000000803 downloaded 03-07-21) protein database that included common contaminants (cRAPome). Searches were performed with settings for the proteolytic enzyme trypsin. Maximum missed cleavages were set to 2. The precursor ion tolerance was set to 10 ppm and the fragment ion tolerance was set to 0.6 Da. Dynamic peptide modifications included oxidation (+15.995 Da on M). Dynamic modifications on the protein terminus included acetyl (+42.011 Da on N-terminus), Met-loss (−131.040 Da on M) and Met-loss+Acetyl (−89.030 Da on M). Static modifications included TMTpro (+304.207 Da on any N-terminus), TMTpro (+304.207 Da on K) and carbamidomethyl (+57.021 on C). Sequest HT was used for database searching. All search results were run through Percolator for scoring and the false discovery rate was set to 1% at the peptide level. Raw quantitative results were transformed to log_2_ scale and normalized to the median value across samples. Missing data were imputed with half of the global minimum value. P-values for pairwise comparisons were calculated by t-test. Datasets are available through MassIVE, accession numbers MSV000094996, MSV000094997, MSV000094998, MSV000094999, MSV000095000, and MSV000095001.

### Publicly available dataset analysis

For publicly available RNAseq and mass spectrometry data from adult fly tissues and staged embryos, data was accessed from Flybase (release FB2024_02) (Öztürk-Çolak et al. 2024).

## ACKNOWLEDGEMENTS

We thank Dr. Ning Zhao and her lab for use of their microscope; the Oliver lab for use of the anti-Tj antibody; C. Smibert for helpful instruction in embryo collection; the Rissland lab for helpful discussion and review; and S. Jagannathan, S. Ramachandran, and S. Nachtergaele for scientific and writing input. This work was funded by NIH grants 5T32GM136444 (C.A.B.), 5T32GM141742 (A.B.R.), R35GM128680 (O.S.R.), and by NSF grant CAREER 2056136 (O.S.R). The Fred Hutch Proteomics and Metabolomics Shared Resource, RRID:SCR_022618, is supported by the Fred Hutch/University of Washington/Seattle Children’s Cancer Consortium (P30 CA015704). The Orbitrap Ascend mass spectrometer used in this research was funded by a grant from the NIH Office of Research Infrastructure Programs (S10 OD030225). This research was supported in part by the Intramural Research Program of the NIH, The National Institute of Diabetes and Digestive and Kidney Diseases (NIDDK) (awarded to Brian Oliver). Some reagents were supplied through the Drosophila Genomics Resource Center, supported by NIH grant 2P40OD010949.

## AUTHOR CONTRIBUTIONS

Conceptualization: O.S.R.; project guidance: O.S.R., P.R.G.; molecular biology and analysis: C.A.B., J.C.H., L.B., and R.G.; imaging and analysis: C.A.B., J.C.H. and A.B.R.; mass spectrometry sample preparation: C.A.B., J.C.H., L.A.J., C.L., P.R.G.; mass spectrometry data analysis: L.A.J., C.L., C.A.B., P.R.G.; writing: C.A.B., J.C.H., and O.S.R.; funding: P.R.G., O.S.R.

## SUPPLEMENTAL TABLES

**Table S1.** Houki immunoprecipitation from ME31B-GFP 0-1 hour embryos mass spectrometry.

**Table S2.** Muskelin-GFP immunoprecipitation mass spectrometry combined with Houki immunoprecipitation mass spectrometry.

**Table S3.** RanBPM knockdown 0-1 hour embryo Muskelin immunoprecipitation mass spectrometry.

**Table S4.** Denaturing crosslinking Muskelin immunoprecipitation mass spectrometry.

**Table S5.** Denaturing crosslinking ME31B immunoprecipitation mass spectrometry.

**Table S6.** Muskelin knockdown 0-1 hour and 3-4 hour embryo whole proteome mass spectrometry combined with RanBPM knockdown 0-1 hour and 4-5 hour whole proteome mass spectrometry.

**Table S7.** Oligonucleotides used in this study.

## SUPPLEMENTAL FIGURE LEGENDS

**Figure S1.** A. Gene name comparisons from yeast, human, and fly CTLH complex homologues. B. Antibody validation. Protein targets were depleted by RNAi in S2 cells followed by western blotting for the target using newly generated antibodies. Gene names above the blots indicate RNAi target, gene names next to blot indicate western blot target. PABP is used as a loading control.

**Figure S2.** A. qPCR confirmation of *Muskelin* mRNA expression in ovary, head, and carcass, normalized to *Muskelin* expression in ovary. mRNA was extracted from each tissue, converted to cDNA, and amplified using gene-specific primers. Expression levels were compared with a student’s t-test, n=3. B, C. Embryo protein (B) and mRNA (C) expression of CTLH components and targets is highest early in development. Each gene was normalized to its maximum expression at each timepoint. D. Muskelin immunofluorescence signal is specific to Muskelin expression. Immunofluorescence was performed on Muskelin-depleted ovaries that express ME31B-GFP E. *Kdo* gene level read coverage tracks for OVO ChIP minus input, GSC ATAC-seq, and *ovo^ΔBP^*/*ovo^ovo-GAL4^*; *UASp-3xFHA-OVO-B* minus *ovo^ΔBP^*/*ovo^ovo-GAL4^*; *UASp-GFP* RNA-seq. Red and black rectangles represent significant OVO DNA binding motifs and OVO ChIP peaks, respectively. Gene models are represented at bottom. Small rectangles represent untranslated regions, large rectangles represent translated regions. Arrows indicate transcriptional start sites.

**Figure S3.** A. Hou immunoprecipitation with the newly developed antibody enriches for CTLH components. Hou was immunoprecipitated from *w1118* 1-2 hour embryo lysate and probed for CTLH complex components. B. Yippee knockdown does not stabilize ME31B-GFP. Embryo lysates were collected across the MZT time course from control embryos or embryos lacking Yippee and probed for ME31B-GFP and actin (as a loading control). C. No putative substrate adaptors are identifiable from comparing Hou immunoprecipitations to Muskelin-GFP immunoprecipitations. Hou immunoprecipitation fold changes compared to control immunoprecipitation were compared to Muskelin-GFP over control fold changes from Cao et al, 2020. In all plots, blue points represent CTLH complex components, orange points represent targets, green points represent bait protein. Each point represents the average fold change of a specific gene across n = 3 replicates. D. Denaturing crosslinking ME31B immunoprecipitation mass spectrometry comparing crosslinked ME31B immunoprecipitation to crosslinked control immunoprecipitation demonstrates CTLH component and repressor complex components as close-proximity binding partners of ME31B in 0-1 hour *w1118* embryo lysate. All western blots are representative of three biological replicates.

